# Directional DBS Leads Show Large Deviations from their Intended Implantation Orientation

**DOI:** 10.1101/631325

**Authors:** TA Dembek, M Hoevels, A Hellerbach, A Horn, JN Petry-Schmelzer, J Borggrefe, J Wirths, HS Dafsari, MT Barbe, V Visser-Vandewalle, H Treuer

**Affiliations:** Department of Neurology, University Hospital of Cologne, Germany; Department of Stereotactic and Functional Neurosurgery, University Hospital of Cologne, Germany; Movement Disorders & Neuromodulation Unit, Department for Neurology, Charité – University Medicine Berlin, Germany; Division of Neuroradiology, Institute of Diagnostic and Interventional Radiology, University Hospital of Cologne, Germany; National Parkinson Foundation International Centre of Excellence, King’s College London, United Kingdom

**Author notes:** contributed equally (TA Dembek, M Hoevels). Corresponding author: Till A. Dembek, Department of Neurology, University Hospital Cologne, Germany, Kerpener Strasse 62, D-50937 Cologne, Germany. Mauritius Hoevels –, Alexandra Hellerbach –, Andreas Horn –, Jan Niklas Petry-Schmelzer –, Jan Borggrefe –, Jochen Wirths –, Haidar Salimi-Dafsari –, Michael T. Barbe –, Veerle Visser-Vandewalle –, Harald Treuer –. Declarations of interest: None. Funding: This investigator-initiated trial did not receive additional funding.

**Keywords:** Deep Brain Stimulation, Directional Stimulation, Lead Orientation, Postoperative CT

## Abstract

**Objective:** Lead orientation is a new degree of freedom with directional deep brain stimulation (DBS) leads. We investigated how prevalent deviations from the intended implantation direction are in a large patient cohort.

**Methods:** The Directional Orientation Detection (DiODe) algorithm to determine lead orientation from postoperative CT scans was implemented into the open-source Lead-DBS toolbox. Lead orientation was analysed in 100 consecutive patients (198 leads). Different anatomical targets and intraoperative setups were compared.

**Results:** Deviations of up to 90° from the intended implantation direction were observed. Deviations of more than 30° were seen in 42 % of the leads and deviations of more than 60° in about 11 % of the leads. Deviations were independent from the neuroanatomical target and the stereotactic frame but increased depending on which microdrive was used.

**Discussion:** Our results indicate that large deviations from the intended implantation direction are a common phenomenon in directional leads. Postoperative determination of lead orientation is thus mandatory for investigating directional DBS.

**Full Disclosures:** Till A. Dembek reports speaker honoraria from Medtronic and Boston Scientific.

Mauritius Hoevels has nothing to disclose.

Alexandra Hellerbach has nothing to disclose.

Andreas Horn reports speaker honoraria from Medtronic.

Jan Niklas Petry-Schmelzer reports travel grants from Boston Scientific.

Jan Borggrefe has nothing to disclose.

Jochen Wirths reports travel grants from Boston Scientific.

Haidar Salimi-Dafsari reports honoraria from Medtronic and Boston Scientific.

Michael T. Barbe reports grants from Medtronic and Boston Scientific.

Veerle Visser-Vandewalle is a member of the advisory boards and reports consultancies for Medtronic, Boston Scientific and St Jude Medical. She received a grant from SAPIENS Steering Brain Stimulation. Harald Treuer has nothing to disclose.

## Introduction

Directional leads have been the latest technological advance in deep brain stimulation (DBS) devices and are now used in the treatment of advanced Parkinson’s disease, essential tremor, and dystonia. Current directional leads feature two electrode levels with three directional electrode segments each to allow axial stimulation steering (Figure 1a). First prospective studies with these leads demonstrated an increase in side-effect thresholds [1] and/or a decrease in efficacy thresholds [2] when using directional stimulation. On the other hand, these new stimulation capabilities come with an increase in programming complexity. Detailed knowledge about the lead’s position with respect to the surrounding anatomy is an important basis for the clinical interpretation of DBS responses and can guide clinicians in selecting the optimal stimulation parameters. In directional DBS however, the orientation angle of the lead now adds a new degree of freedom which has to be known to relate stimulation parameters to neuroanatomy. While neurosurgeons typically intend the lead to face into a certain direction, it is unclear whether this can be achieved with the current intraoperative techniques. Therefore, a reliable method to determine lead orientation from postoperative imaging is needed. We thus developed and released the Directional Orientation Detection (DiODe) algorithm which calculates the orientation of directional DBS leads based on postoperative CT-scans [3–5]. In this study, we aimed to investigate whether deviations from the intended implantation direction occur by analyzing the orientation of directional DBS leads in a large and heterogeneous patient cohort.

**Figure 1.**
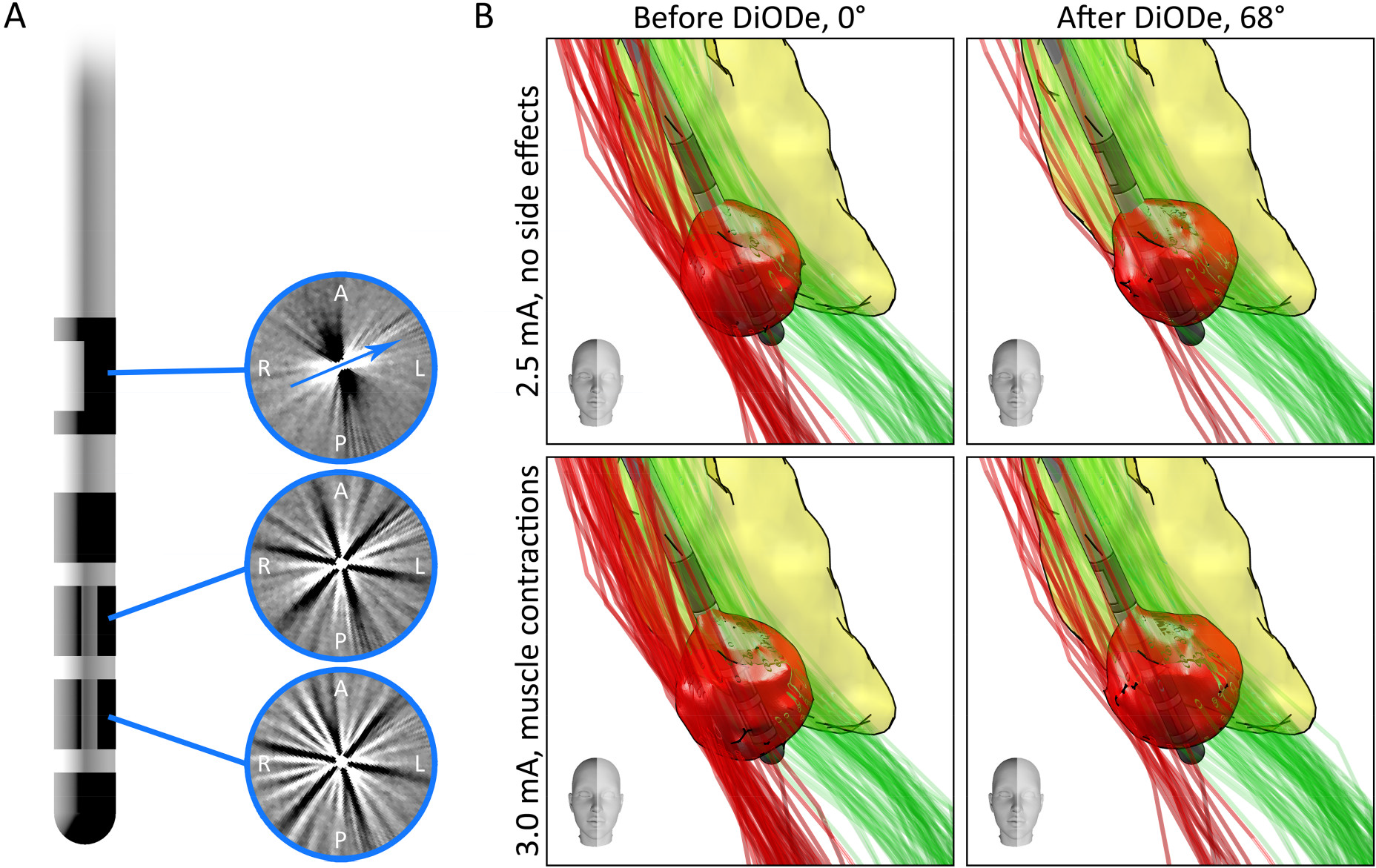
DiODe. A) CT artifacts generated by the stereotactic marker and the segmented electrodes. The blue arrow indicates the orientation of the lead. B) Example application in a patient with essential tremor. The right lead is shown from anterior with the intended orientation angle of 0° (left) and with the real orientation angle of 68° (right). VTAs are shown for stimulation on electrode 13 and amplitudes of 2.5 mA (above) and 3.0 mA (below). Fibers activated by stimulation of a connectome (Horn et al. 2014 [3]) based dentato-rubro-thalamic-tract (DRTT) are shown in green, while activated fibers belonging to the cortico-spinal-tract (CST) are shown in red. The premotor thalamus is shown in yellow [12]. At 2.5 mA the patient experienced full tremor control and no observable side-effects, while muscle contractions indicating an affection of the CST began at 3.0 mA. Ignoring the real lead orientation led to widespread activation of the CST in both settings while incorporating the orientation into the analysis only showed CST activation at 3.0 mA, when side-effects were present.

## Methods

### Clinical data

We retrospectively analyzed data from the first 100 consecutive patients who had been implanted with Cartesia™ directional DBS leads (Boston Scientific, USA) in our center. Patients had been selected for DBS treatment according to our clinical routine. The cohort included different neurological indications and surgical targets (see results). Due to the retrospective nature of our analysis no local ethics approval or written informed consents were obtained.

### Implantation and imaging

All patients received preoperative MRI and CT imaging for stereotactic planning. Two neurosurgeons performed the implantations, using different stereotactic systems (CRW^®^ frame, Integra Life Sciences, USA; RM^®^ frame, Inomed, Germany) respectively. The stereotactic marker was implanted to face anteriorly (orientation angle of 0°) which was controlled visually via either intraoperative fluoroscopy (CRW) or intraoperative stereotactic x-ray (RM) before fixation of the lead. During the course of the study, the intraoperatively used microdrives changed from an ISIS MER system (Inomed, Germany) to a Neuro Omega™ system (AlphaOmega, Israel). Patients received postoperative CT imaging as part of their routine clinical care to exclude perioperative bleeding and to confirm correct lead positioning. CT scanners iCT256 and IQon (Philips, The Netherlands) were used with the following scan parameters: pixel size 0.6 mm (range: 0.4 – 0.7 mm), slice thickness 0.8 mm (range: 0.6 – 1.0 mm), spiral pitch factor 0.36 (range: 0.34 – 0.44), gantry tilt 0°, tube voltage 120 kV, exposure 240 mAs, filter type UB (soft tissue).

### Orientation detection from postoperative CT

The DiODe algorithm is based on previous works from our center and the detailed methodology is explained elsewhere [4,5]. In short DiODe inspects the artifacts generated by the Cartesia leads (Figure 1a) in postoperative CT scans to determine the orientation of the lead. In a first step, the orientation of the artifact generated by the stereotactic marker is analyzed. Afterwards the characteristic streak artifacts generated by the directional electrode segments are used to calculate a more accurate estimation. The original user-supervised algorithm, which was implemented in IDL (Exelis Visual Information Solution, USA), has been extensively validated in both geometrical and anthropomorphic phantoms [4,5]. It yielded accurate and robust results, as long as the polar angle between the lead and the CT scanner axis remained below 60°. We now provide an adaptation of DiODe implemented in Matlab (The MathWorks, USA) and fully integrated into the open-source Lead-DBS toolbox (https://www.lead-dbs.org) [3]. Lead-DBS enables the user to a) conduct coregistration of preoperative and postoperative imaging, b) to normalize images to different atlas spaces, and c) to reconstruct lead trajectories. Based on these trajectories DiODe automatically extracts the relevant artifacts and calculates the orientation angles in a matter of seconds (*automatic* workflow). The user is then required to visually inspect and confirm the results or, if deemed necessary, to use a *manual refine* workflow to improve results by respecifying the CT slices where the artifacts are most visible, respecifying the centers of the artifacts within the slices, and/or by choosing a different solution for the symmetric marker artifact. The results are automatically propagated from the CT space into both the patient MRI and the atlas space. The Matlab implementation of DiODe was extensively validated against the previously published version (see Data Supplement). The *automatic* workflow yielded accurate results for polar angles < 40° while the *manual refine* workflow was accurate for polar angles < 55°.

### Image analysis in Lead-DBS

Preoperative MRI and postoperative CT images were coregistered linearly using Advanced Normalization Tools (ANTs) in all patients [6]. Images were then normalized to standard stereotactic space (MNI ICBM 2009b, asymmetric) using nonlinear ANTs with subcortical refine [6–8]. Lead trajectories were identified using the PaCER algorithm [9]. Afterwards the DiODe algorithm was used to determine the orientation angles. In a first step, only the *automatic* workflow was used. In a second step, results were inspected and, if warranted, improved via the *manual refine* workflow. Both workflows were used for validation (see Data Supplement).

### Analysis of deviations

The final orientation results after manual refinement were used to investigate deviations from the intended implantation orientation (anterior, 0°) with respect to the patient’s coordinate system defined by the line connecting anterior commissure (AC) and posterior commissure (PC). The prevalence of deviations was explored using histograms. To test whether there was a systematic deviation from 0° we used one-sample Wilcoxon signed-rank test with a significance threshold of p < 0.05. We also investigated differences in deviations for different anatomical targets using the nonparametric Kruskal-Wallis test and post-hoc Wilcoxon rank-sum test with significance thresholds of p < 0.05, Bonferroni-corrected for multiple comparisons. Additionally, we compared our different intraoperative setups (CRW-frame, intraoperative fluoroscopy versus RM frame, intraoperative stereotactic x-ray) and the two microdrives (AlphaOmega versus Inomed) to see whether deviations differed between approaches (Wilcoxon rank-sum test, p < 0.05). Furthermore differences in variance were analyzed using the Brown-Forsythe test (p < 0.05).

## Results

### Data

One-hundred patients receiving a total of 198 directional leads were analyzed. Surgical targets were the subthalamic nucleus (STN) in 101 leads, the ventral intermediate nucleus and the posterior subthalamic area (VIM/PSA) in 69 leads, and the internal part of the globus pallidus (GPI) in 28 leads. The CRW frame was used in 98 leads while the RM frame was used in 100 leads. The Inomed system was used in 115 leads, while the AlphaOmega system was used in 83 leads.

### Prevalence of deviations

Large deviations ranging from −89.1° to 88.1° with respect to the desired implantation direction of 0° were observed in our cohort as can be seen in Figure 2. Deviations of more than 30° occurred in 82 leads (41 %) and deviations of more than 60° in 23 leads (11 %). The median deviation was −4.3° (interquartile range: −26.7° to 21.5°) and did not differ from 0° (p = 0.22), indicating that leads did not systematically deviate into one direction. No differences in deviations (p = 0.73) or differences in variances (p = 0.26) were observed for the different anatomical targets STN, PSA/VIM, and GPI. Also there was no difference regarding deviation (p = 0.46) and variance (p = 0.53) between the two different intraoperative setups (frame, surgeon, intraoperative imaging). While deviations did not differ when comparing the two microdrives (Inomed versus AlphaOmega, p = 0.59) variance was significantly increased by a factor of 2.24 when using the AlphaOmega system (p < 0.0001, Figure 2).

**Figure 2.**
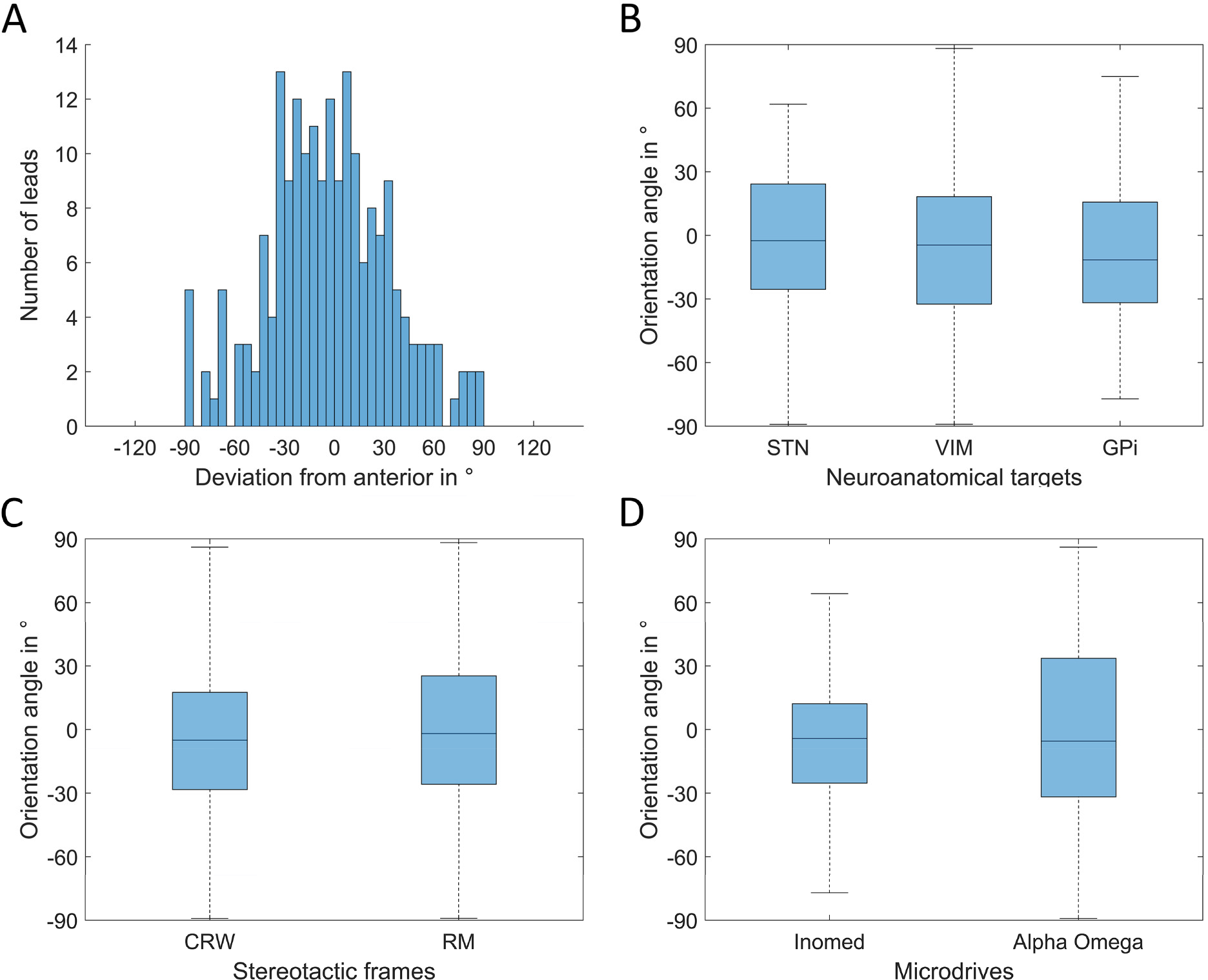
Results. A) Histogram of orientation angles with respect to the intended orientation (anterior, 0°). B) Boxplot depicting the orientation of leads implanted into the STN, the VIM, and the GPI. C) Boxplot depicting the orientation of leads implanted using either the CRW or the RM stereotactic frame. D) Boxplot depicting the orientation of leads implanted using either the Inomed or the AlphaOmega microdrive. Variance was increased with the AlphaOmega compared to the Inomed system (p < 0.0001).

## Discussion

This is the first study investigating the orientation of directional DBS leads in patients. In this large cohort of 100 patients, deviations of up to 90° from the intended implantation direction could be observed. In more than 10 % of the leads the deviation was more than 60°. This was independent from the surgical target and the intraoperative setting comprising of stereotactic frame, neurosurgeon, and intraoperative imaging modality. However, the variance increased depending on the intraoperatively used microdrive. The prevalence and amount of deviations might sound surprising at first, but many factors may impede the accurate placement of the lead into a predefined direction. First and foremost, while the depth of the lead can be adjusted using a microdrive, the orientation of the lead has to be adjusted manually. Considering the small diameter of the lead, a manual correction by 10° for example would require the neurosurgeon to turn the lead by just 0.1 mm, which is almost impossible to achieve by hand. Furthermore, while the neurosurgeon intends to position the lead into a certain direction with respect to the patient’s neuroanatomy, the neuroanatomic frame of reference is elusive in most intraoperative settings. Instead the neurosurgeon has to use detached landmarks to guide manual rotation (e.g. skull features). Finally, the lead in itself is not stiff but susceptible to internal torsion [10], some of which may occur during or even after lead fixation. Concluding that accurate manual orientation of directional DBS leads is not possible in a relevant amount of cases, a reliable way to determine the lead orientation postoperatively gains even more importance.

A few other approaches have been proposed to determine the orientation of directional leads from intraoperative stereotactic x-ray, rotational fluoroscopy, or flat-panel CT [4,10,11]. The approach used in this study has the advantage, that postoperative CT imaging is already part of the routine care in many centers to verify lead location and to exclude intracerebral hemorrhage. Second, other methods relied on subjective user assessment [10,11]. While those methods are accurate and show high inter-rater reliability, we think that an automatic approach is less cumbersome and increases comparability. Our algorithm is able to provide accurate results in a few seconds and even the manual refine workflow takes less than a minute – much less than user-dependent approaches. Third, our algorithm in combination with Lead-DBS automatically translates the lead’s orientation into a variety of neuroanatomical spaces. This allows users to investigate lead orientation in relation to individual anatomy, neuroanatomical atlases, tractography and connectomes, and to use simulated stimulation volumes (VTAs) to investigate the origins of directional DBS effects (Figure 1b).

Concluding, we demonstrate that large deviations from the intended implantation direction are seen in a significant proportion of patients. To address this we provide a freely-available algorithm which can reliably detect the orientation of directional DBS leads from postoperative CT with only minimal user-intervention.

## Contributions

TAD – Study conception, development of methods, data & statistical analysis, drafting of the manuscript

MH – Study conception, development of methods, data analysis, critical revision of the manuscript

HA – Study conception, development of methods, data analysis, critical revision of the manuscript

AH – Development of methods, critical revision of the manuscript

JNPS – Development of methods, critical revision of the manuscript

JB – Development of methods, critical revision of the manuscript

JW – Data acquisition, critical revision of the manuscript

HSD – Data acquisition, critical revision of the manuscript

MTB – Data acquisition, critical revision of the manuscript

VVV – Study conception, data acquisition, critical revision of the manuscript

HT – Study conception, development of methods, data acquisition & analysis, drafting of the manuscript

All Authors gave final approval of this manuscript to be submitted.

## Data Supplement - Validation of the DiODe algorithm

### Introduction

The original user-supervised version of the DiODe algorithm, was implemented in IDL (Exelis Visual Information Solution, USA), and has already been extensively validated in both geometrical and anthropomorphic phantoms.^1,2^ In this data supplement we now validate both the *automatic* and the *manual refine* workflows of the Matlab-based version as it was implemented into the Lead-DBS toolbox.

### Data

Lead orientation was analyzed in all 198 leads as described in the methods section of the main article using a) the *automatic* workflow. All results were then inspected and b) the *manual refine* workflow was used when an experienced developer of the algorithm (TAD) deemed that optimization was possible. To validate our results we used two independent methods: c) the IDL-based, user-supervised algorithm^2^ in all leads and d) a previously published method for intraoperative stereotactic x-ray^1^ in all leads where stereotactic x-ray was available (n = 100).

### Analysis

Results from the automatic and the manual refine workflow were correlated to the results from the IDL-based, user-supervised algorithm and to results from stereotactic x-ray using nonparametric Spearman-correlations. Deviations between results were described using histograms and by calculating the median as well as the interquartile range. Additionally we examined all individual cases in which the deviation between the Matlab and the IDL-based algorithm was larger than 10°.

### Results

All 198 leads were analyzed regarding their orientation using the DiODe *automatic* workflow. After visual inspection of the results, the experienced user tried the *manual refine* workflow in 137 leads to see whether optimization was possible.

### Validation against IDL-based algorithm for postoperative CT

Spearman correlation revealed highly significant and strong correlations between the results of the IDL-based algorithm and the DiODe *automatic* workflow (rho = 0.80, p < 0.0001). The histogram revealed that when using the *automatic* workflow orientations in 182 leads (92 %) deviated less than 10° from the IDL-based algorithm (176 leads (89 %) less than 5°). The median deviation was −0.6° with an interquartile range of −1.8° to +0.6°. In 137 out of 198 leads the user decided to try further manual refinement by either respecifying the analyzed CT slices, the exact lead position, or the solution chosen by the algorithm. However the resulting orientation angles of the *manual refine* workflow only differed by more than 10° from the results of the *automatic* workflow in 14 of the 137 cases (10 %). When comparing results after *manual refinement* to the IDL-based algorithm, Spearman correlation again revealed highly significant and strong correlations (rho = 0.90, p < 0.0001). After *manual refinement* 191 leads (96 %) deviated less than 10° from the IDL-based algorithm (188 leads (95 %) less than 5°). The median deviation was −0.6° with an interquartile range of −1.4° to +0.2° (Figure 1 a-c).

### Posthoc analysis of large deviations

In a post-hoc analysis we investigated all cases which showed deviations of larger than 10° (*automatic* n = 16; *manual refine* n = 7) from the results of the IDL-based algorithm. In 5 *automatic* cases and 3 *manual refine* cases the solutions between DiODe and IDL differed by about 180° implying that DiODe chose a different solution to the orientation angle of the symmetric marker artifact than IDL. In all those cases orientation angles deviated by about 90° from the planned anterior direction so that the marker was either facing medially or laterally and the correct solution could not be properly assumed without additional information. In n = 11 *automatic* and n = 4 *manual refine* cases high polar angles of the trajectories with respect to the CT scanner axis were observed. In the 11 *automatic* cases the median polar angle was 49.1° (range 40.9°-66.7°) while it was 59.1° (range 54.8°-66.7°) in the remaining 4 *manual refine* cases. The n = 2 remaining *automatic* cases only differed from the IDL solution by 12.4° and 20.9° (Figure 2).

### Validation against algorithm for stereotactic x-ray

Stereotactic x-ray could be successfully analyzed in 90 of 100 leads. Results of both the *automatic* and the *manual refine* workflow correlated strongly with results from stereotactic x-ray (*automatic*: rho = 0.83, p < 0.0001; *manual refine*: rho = 0.94, p < 0.0001). The median deviation was −1.1° (interquartile range −7.5° to +6.5°) for the automatic and −2.5° (interquartile range −7.6° to +4.5°) for the manual refine workflow respectively (Figure 1 d-f).

### Interpretation

This validation provides evidence that the DiODe algorithm is able to reliably and accurately detect the orientation in the vast majority of cases. Two independent measures, both extensively validated in phantoms, were used as gold standards providing reliability of the results. The *automatic* workflow provided accurate results as long as the polar angle between the lead and the CT scanner axis was less than 40°. In cases with larger polar angles, the *manual refine* workflow provided accurate results for polar angles up to 55°. In conclusion, two recommendations can be given to users of the algorithm: First, large polar angles should be avoided. This can be achieved mainly by correct positioning of the patient inside the CT scanner. Additional flexion of the neck (e.g. by using a cushion) should be avoided, because it increases the polar angle. Instead patients should be positioned flatly on the CT scanner table. If one likes to avoid radiation exposure of the eyes, we recommend using protective eye covers which in our experience do not impede the results of the algorithm. Second, in those cases were a large deviation of the intended implantation direction is seen, we recommend performing an additional single plane x-ray. This can ensure that the algorithm picks the right solution in those cases where it is unclear whether the lead orientation deviates 90° to the left or 90° to the right.

**Figure 1.**
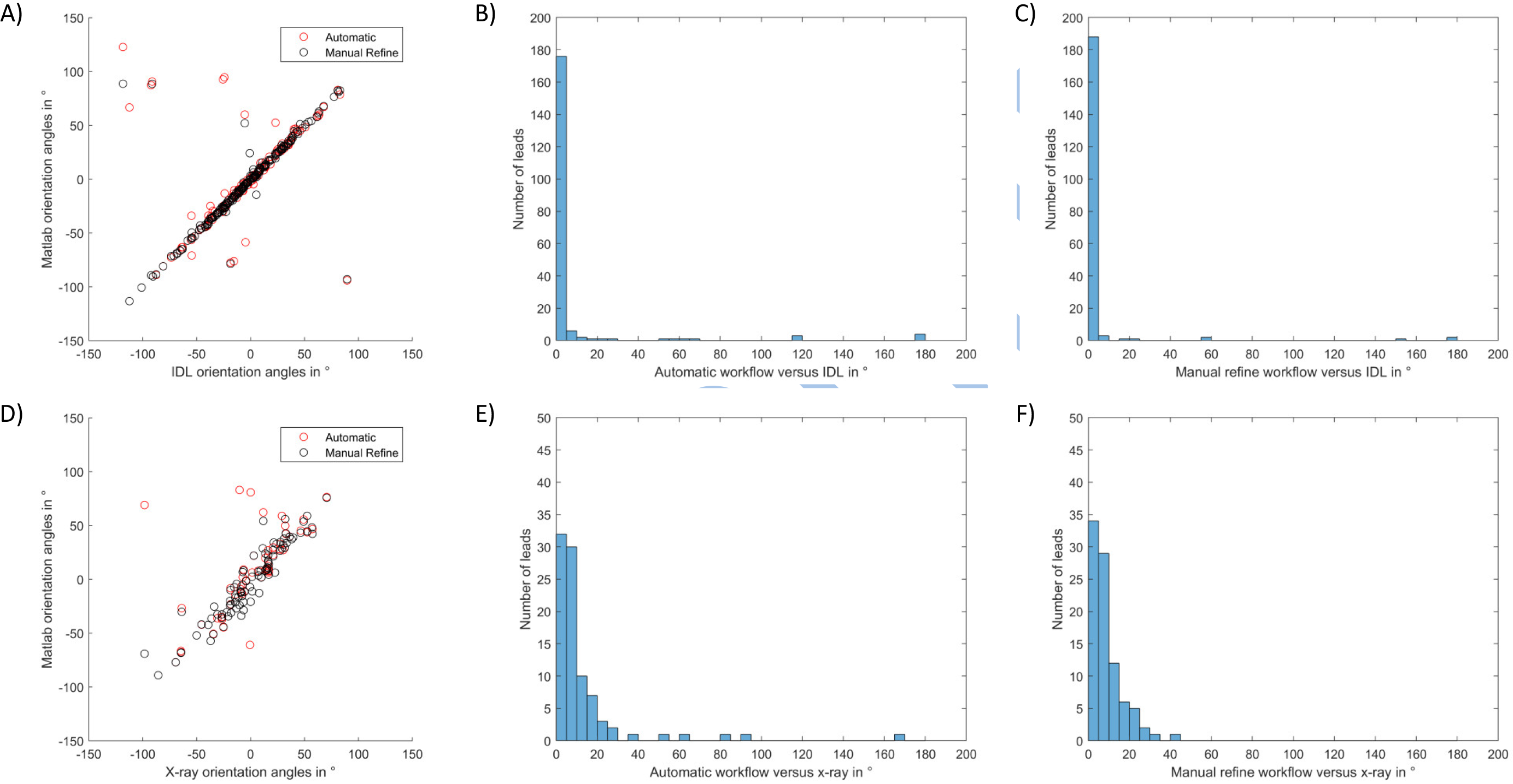
Results. Validation of the DiODe automatic and manual refine workflows against the IDL-based algorithm for postoperative CT (A-C) and against stereotactic x-ray (D-F). A) Scatter plot of orientation angles generated by the automatic workflow (red) and the manual refine workflow (black) compared to orientation angles generated by the IDL-based algorithm. B+C) Histograms of the differences in orientation angles between the IDL-based algorithm and the automatic/manual refine workflows respectively. D) Scatter plot of orientation angles generated by the automatic workflow (red) and the manual refine workflow (black) compared to orientation angles generated from stereotactic x-ray. E+F) Histograms of the differences in orientation angles between stereotactic x-ray and the automatic/manual refine workflows respectively.

**Figure 2.**
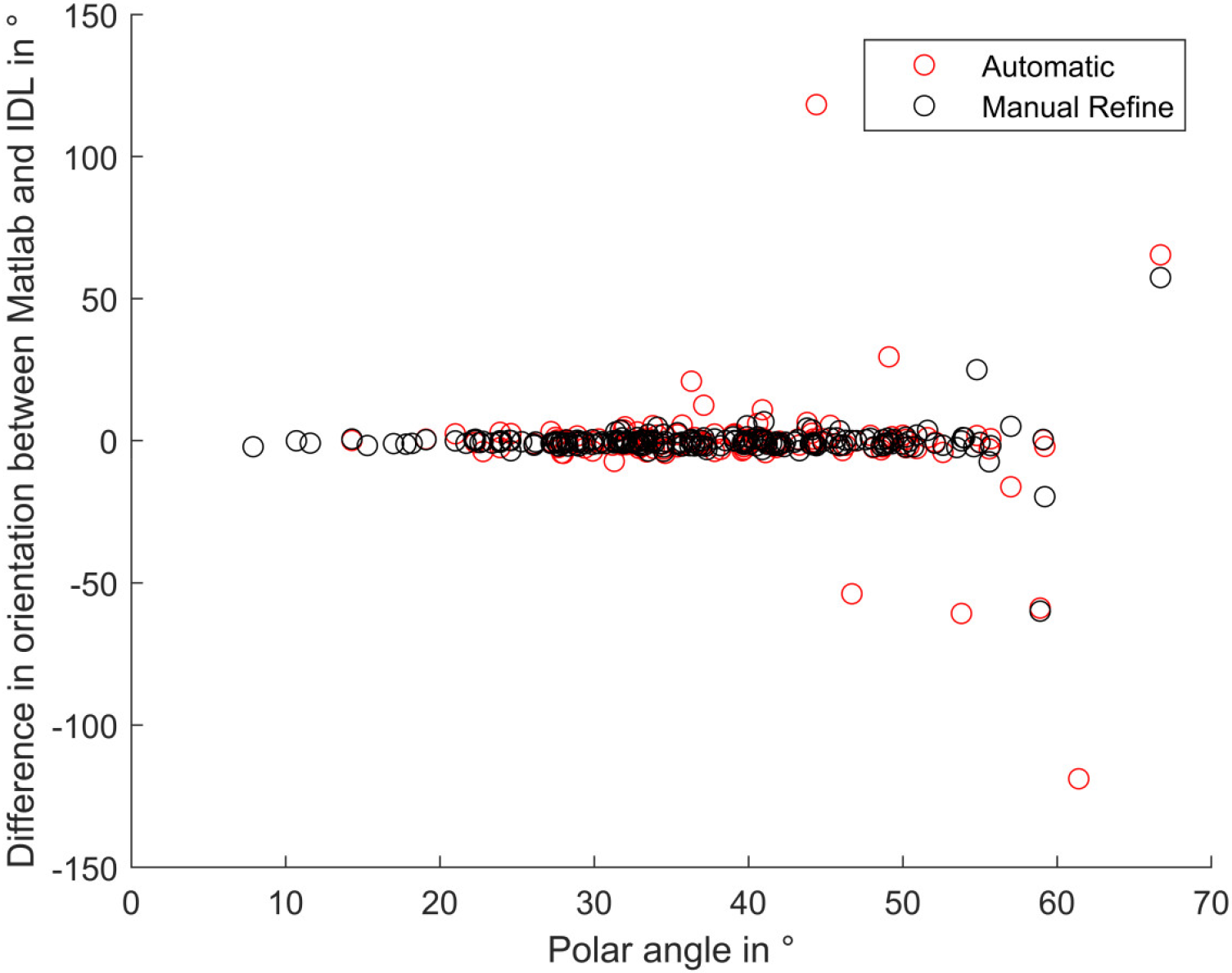
Dependency from Polar Angle. Orientation differences between the IDL-based algorithm and the *automatic* (red) and *manual refine* (black) workflows with relation to the polar angle of the lead’s trajectory. After *manual refinement*, large differences were only observed for polar angles > 55°, while for the *automatic* workflow they were only seen for polar angles > 40°.

## Notes

https://www.lead-dbs.org

